# Hyperactivation of Mincle in exacerbated apical periodontitis in an autoinflammatory mouse model for cherubism

**DOI:** 10.64898/2026.11.09.765482

**Authors:** Ayano Hatori, Kiseki Washitani, Eisaku Nishida, Daniel Faissal Kotait, Leslie Blumenfeld, Yasuyoshi Ueki, Ernst J Reichenberger, Adam Kim, I-Ping Chen

## Abstract

**Aims:** To examine the role of Mincle, a C-type lectin (*Clec*) pathogen recognition receptor (PRR), in exacerbated apical periodontitis (AP) in an autoinflammatory mouse model for cherubism (CBM).

**Methodology:** Patients with immune system dysfunction have higher prevalence of AP but the mechanism remains largely unknown. We studied a knock-in (KI) mouse model carrying a *Sh3bp2* mutation for CBM. We performed pulp exposure in one lower 1^st^ molar of 5- to 6-week-old mice to introduce pathogen-associated molecular patterns (PAMP)-induced endodontic infection. We determined the expression levels of *Clec* subfamily genes by qPCR. To suppress Mincle, we generated *Sh3bp2^+/+^Mincle^+/+^*, *Sh3bp2^+/KI^Mincle^+/+^*, and *Sh3bp2^+/KI^Mincle^KO/KO^* mice or injected *Sh3bp2^+/KI^* mice with a Mincle-binding DNA aptamer (AptMincle). Mice were analyzed 42 days after pulp exposure by micro-CT (μCT), qPCR and immunoblotting. Statistical analysis was performed by one-way ANOVA followed by post-hoc analysis with Tukey’s test or two-way ANOVA followed by Sidak correction. *p* < 0.05 was considered statistically significant.

**Results:** Expression of *Clec4e* (*Mincle*) was significantly increased in *Sh3bp2* mutant mice after pulp exposure compared to WT littermates (*p*<0.0001). *Sh3bp2^+/KI^Mincle^+/+^* and *Sh3bp2^+/KI^Mincle^KO/KO^* mice had a comparable response to pulp exposure, including the degree of bone loss and induction of inducible *NOS (iNOS)*, *IL-1b*, and *TNF-a*. *Sh3bp2^+/KI^* mice injected with AptMincle after pulp exposure for 21 days showed significant amelioration of bone loss compared to mice with vehicle injection (*p*<0.0001).

**Conclusions:** These seemingly conflicting data of suppressing Mincle by crossing with Mincle KO mice and AptMincle injection suggest that Mincle has dual functions upon PAMPs stimulation, consistent with a previous study showing that Mincle enhances tissue destruction during the inflammatory stage and is subsequently needed for inflammation resolution. Fine-tuned suppression of Mincle significantly reduces the severity of AP in *Sh3bp2^+/KI^* mice. This study suggests that patients with CBM may benefit from more frequent dental examinations and early intervention to reduce or prevent severe progressive bone loss in periapical tissues.

## Introduction

Apical periodontitis (AP) is primarily caused by microbial invasion of the root canal system and periapical tissues, which stimulates the host immune system to produce inflammatory responses (1). Patients with immune system dysfunction may develop excessive tissue destruction in AP. The prevalence of AP in patients with inflammatory bowel disease, rheumatoid arthritis, and psoriasis, is significantly higher than in unaffected controls, likely due to increased infection susceptibility and impaired healing mechanisms (2, 3). Moreover, patients with inflammatory bowel disease develop significantly larger lesions than healthy subjects (4). How host immunity affects the development of AP is not fully understood, in part due to limited case access. Here, we examine AP development in a mouse model for cherubism (CBM). Mutant mice represent an autoinflammatory disease model because their systemic hyperinflammation is T/B cell independent and primarily driven by myeloid cells (macrophages and osteoclasts) (5, 6).

CBM is a rare genetic autoinflammatory bone disorder, characterized by expansile jawbones with multilocular fibro-cystic lesions (7, 8). Gain-of-function mutations in Src Homology 3 Domain-Binding Protein 2 (*SH3BP2;* mouse orthologue *Sh3bp2*), an adaptor protein involved in immune cell development and immune response, have been identified for the autosomal dominant CBM (9). Knock-in (KI) mice carrying a homozygous *Sh3bp2_P416R_* mutation (*Sh3bp2^KI/KI^*) develop trabecular bone loss, cortical bone erosion, and systemic hyperinflammation with elevated TNF-α expression. *Sh3bp2^KI/KI^* mice have increased Erk1/2 and Syk phosphorylation/activation in myeloid cells (5, 10–13). Although CBM is an autosomal dominant disease, *Sh3bp2^+/KI^* mice exhibit an intermediate phenotype, resembling WT mice in gross appearance (5, 14). We have previously induced AP by exposing pulp in one mandibular molar of *Sh3bp2*^+/+^, *Sh3bp2^+/KI^*, and *Sh3bp2^KI/KI^* mice. Compared to *Sh3bp2^+/+^* mice, where apical lesions were smaller in size and localized, *Sh3bp2^+/KI^* and *Sh3bp2^KI/KI^* mice develop exacerbated AP with increased bone loss, expansile jawbones, and hyperactivated neutrophils, macrophages, and osteoclasts (15).

Pathogen-associated and damage-associated molecular patterns (PAMPs & DAMPs) are recognized by pattern recognition receptors (PRRs), such as Toll-like receptors (TLRs), C-type lectin receptors (CLRs, *Clec* family genes), nucleotide-binding oligomerization domain-like receptors (NODs), and retinoic acid-inducible gene (RIG) 1-like receptors (16, 17). TLR2 and TLR4 are the most well-studied PRRs contributing to AP, by activating immune cells to secrete proinflammatory mediators, promoting angiogenesis, and by enhancing synthesis of reactive oxygen species (18, 19). We previously investigated *Tlr2/4*-mediated hyperinflammation in CBM and showed that *Sh3bp2^KI/KI^Tlr2/4^KO/KO^* mice were protected from jawbone expansion (15). Since we suspected that other PRRs could be involved in endodontic infection of CBM mice we focused on the role of *Clec* genes. CLRs expressed on immune cells of both the innate and adaptive immune systems serve as important regulators in immune response and homeostasis (20). Several types of CLRs, such as Mincle (*Clec4e*, macrophage-inducible C-type lectin), can activate pathways through immunoreceptor tyrosine-based activation motifs (ITAMs) and the adaptor Fc receptor γ-chain (FcRγ) to further facilitate the recruitment of spleen tyrosine kinase (SYK) (21). These intracellular signals primarily result in the activation of nuclear factor-κB (NF-κB)-dependent pro-inflammatory responses. Interestingly, SYK, NF-κB, and NFAT signaling pathways, which are downstream of CLRs, are highly upregulated in *Sh3bp2* mutant mice (5). Here, we focus on CLRs in *Sh3bp2^+/KI^* mice because CBM is an autosomal dominant disease. We hypothesized that *Sh3bp2^+/KI^* mice express significantly more *Clec* genes upon pathogen stimulation, which in turn contributes to exacerbated AP. We propose that C-type lectin receptors are a missing link for hyperinflammatory responses in CBM.

## Materials and Methods

The manuscript of this animal study has been written according to the Preferred Reporting Items for Animal studies in Endodontology (PRIASE) guidelines (22).

### Mice

*Sh3bp2* (P416R) knock-in mice have been previously reported (5). A *Mincle^KO/KO^* mouse model was a gift from Dr. Adam Kim. These mice were originally generated at Osaka University (23). We bred *Sh3bp2^+/KI^Mincle^+/KO^* double mutant mice to obtain control mice (*Sh3bp2^+/+^Mincle^+/+^, Sh3bp2^+/KI^Mincle^+/+^*) and experimental mice (*Sh3bp2^+/KI^Mincle^KO/KO^*). All strains were on C57BL/6J background housed in a specific pathogen-free (SPF) environment with a normally complex microflora. All experiments involving animals were performed in accordance with animal protocol AP-200505-0424 approved by the Animal Care Committee at the University of Connecticut Health (UConn Health).

### Pulp exposure in mice

We performed pulp exposure of left mandibular first molars in 5- to 6-week-old male and female mice using a #1/4 round carbide bur powered by an electric handpiece (AEU-20, Tulsa Dental, Tulsa, USA) under a dissection microscope (Zeiss, Oberkochen, Germany) with isoflurane sedation as described previously (15). Pulp was left open for 42 days in *Sh3bp2^+/KI^* mice or 14 days in *Sh3bp2^KI/KI^* mice to allow for the development of AP. Mice were monitored twice a week to ensure no pain and no disability after pulp exposure.

### AptMincle injection

We adapted an AptMincle *in vivo* injection protocol (24). *Sh3bp2^+/KI^* mice were injected with AptMincle (3.5 mg/kg in a total volume of 200 μL) by intraperitoneal injections on the day of pulp exposure and twice a week for 3 weeks. Vehicle treated *Sh3bp2^+/KI^* mice had pulp exposure but received saline only. All mice were euthanized 42 days after pulp exposure.

### Radiographic and μCT imaging analysis

Radiographs of mandibles were obtained by a KUBTEC radiography system (KUBTEC Medical Imaging, Trumbull, USA). Bony expansion after endodontic infection in *Sh3bp2* mutant mice was measured as previously reported (15). We measured changes in expansile mandibles normalized to the length of roots, which is a parameter that does not change due to pulp exposure. Radiographic analysis was performed with Photoshop CC (Adobe, San Jose, USA). μCT scanning was performed by μCT core facility (mCT20; Scanco Medical, Switzerland) at UConn Health. Destruction of alveolar bone after pulp exposure was measured by bone volume fraction (BVF) (bone volume/total volume; BV/TV%) obtained from 80 sections (12 micron/section) of coronal planes covering centers of mesial and distal roots of the mandibular 1^st^ molars. Incisors were excluded from the measurement.

### RNA isolation and qPCR analysis

We isolated RNA from mouse mandibles and performed qPCR analysis as previously described (15). Briefly, mandibles were dissected out in RNAlater (Millipore Sigma, Burlington, USA) and the first molar with surrounding alveolar bone was immediately frozen in liquid nitrogen. Incisors were excluded. The frozen bone was crushed using a cryogenic tissue pulverizer (Fisher Scientific, Waltham, USA). Bone powder was immersed in TRIzol (Invitrogen, Carlsbad, USA), homogenized (Ultra-Turrax; IKA-WERKE, Wilmington, USA), and centrifuged. RNA isolation was performed using a Direct-zol RNA kit (Zymo Research, Irvine, USA). DNAse treatment was performed during RNA isolation. qPCR using Universal SYBR Green Supermix (BioRad, Hercules, USA) was performed in a CFX connect Real-Time System (BioRad, Hercules, USA). PCR efficiency was optimized, and primer specificity was confirmed by melting curve analysis. Gene expression levels were calculated using the 2^−ΔΔCt^ method and normalized to *18S* RNA as endogenous control. Primer sequences used are shown in S1 Table.

### Immunoblotting

We prepared whole-cell lysates from mandibular bones using RIPA buffer (Tris 50 mM, pH=7.8, NaCl 150 mM, SDS 0.1%, sodium deoxycholate 0.5%, NP40 1%) containing Halt Protease and Phosphatase Inhibitor Cocktail (Thermo Fisher Scientific, Waltham, USA). Mandibular bones were frozen in liquid nitrogen and crushed using a cryogenic tissue pulverizer (Fisher Scientific, Waltham, USA). RIPA lysate was collected by placing samples on a shaker in the cold room, homogenized with a homogenizer (Ultra-Turrax, IKA-Werke, Wilmington, USA), and centrifuged at high speed for 10 minutes. The concentration of protein was determined by BCA protein assay (Thermo Fisher Scientific, Waltham, USA). Protein lysates were loaded onto 12% SDS-PAGE gels. Samples were then transferred onto PVDF membranes using a wet transfer apparatus (BioRad, Hercules, USA). To reduce nonspecific binding, membranes were blocked in 5% skim milk for 1 hour at room temperature. The membranes were incubated overnight with anti-Mincle antibody (1:1000; D292-3; MBL Life Science, Tokyo, Japan) and anti-actin antibody (1:1000; sc-47778, Santa Cruz Biotechnology, Dallas, USA). Primary antibody was diluted in Tris-buffered saline containing 0.1% Tween-20 and 1% skim milk. After incubation with primary antibodies, the membranes were washed in Tris-buffered saline/Tween buffer and then incubated with corresponding secondary antibodies (1:10,000; Thermo Fisher Scientific, Waltham, USA) in Tris-buffered saline/Tween buffer containing 5% skim milk for 1 hour. Protein bands were detected by enhanced chemiluminescent detection reagent (Azure Biosystems, Dublin, USA) and visualized by an Azure c600 imaging system (Azure Biosystems, Dublin, USA).

### Statistics

Statistical analysis was performed by one-way ANOVA followed by post-hoc analysis with Tukey’s test or two-way ANOVA followed by Sidak correction (for multiple comparisons) using Prism 8 software (GraphPad Software, San Diego, USA). *p* < 0.05 was considered statistically significant.

## Results

### Significantly increased *Clec4e* (Mincle) in *Sh3bp2^+/KI^* mice after pulp exposure

CLRs (*Clecs*) are composed of a family of more than 1000 proteins, consisting of 17 groups based on their structure and function (20, 25). We focused on genes in CLR Groups II and V, belonging to Dectin-1 and Dectin-2 clusters, which recognize pathogens and ligands on damaged cells. These CLRs trigger inflammatory pathways via the ITAM-FcRγ-Syk axis. To identify CLRs that are differentially expressed in *Sh3bp2^+/+^* and *Sh3bp2^+/KI^* mice and are induced by pulp exposure, we measured expression of several *Clec genes* in pulp-exposed vs. no-pulp-exposed mandibles by qPCR. We showed that *Clec4e* (*Mincle*), *Clec4d* (*Dectin-3*), and *Clec6a* (*Dectin-2*) were significantly triggered in *Sh3bp2^+/KI^* mice on the sites of pulp exposure compared to the contralateral sides without pulp exposure and to *Sh3bp2^+/+^* mice with or without pulp exposure (Fig 1A). *Clec4a2* (*Dcir*) and *Clec12a* expression levels showed no significant difference between *Sh3bp2^+/+^* and *Sh3bp2^+/KI^* mice with or without pulp exposure (Fig 1B). *Clec2d* was equally induced in *Sh3bp2^+/+^* and *Sh3bp2^+/KI^* mice after pulp exposure (Fig 1B). Among *Clec* genes, *Clec4e* (*Mincle*) was highest expressed and most significantly induced after pulp exposure in *Sh3bp2^+/KI^* mice. We next examined whether Mincle plays an important role in AP development of *Sh3bp2^+/KI^* mice. We hypothesize that suppression of *Clec4e* (Mincle) can partially ameliorate bone loss in AP of *Sh3bp2^+/KI^* mice.

**Fig 1:**
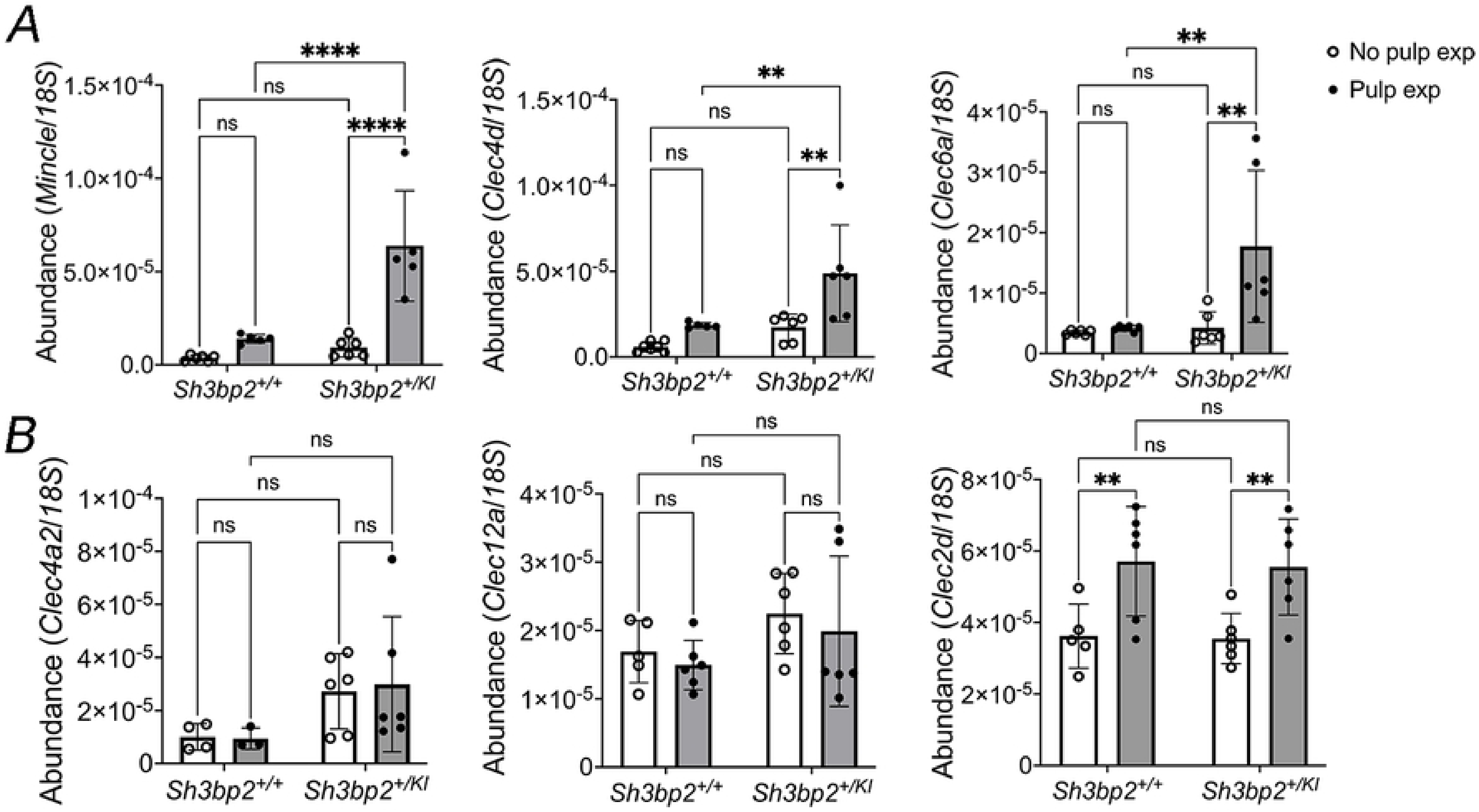
Increased expression levels of Mincle in *Sh3bp2^+/KI^* mice after pulp exposure. qPCR data show **(A)** *Clec4e (Mincle)*, *Clec4d,* and *Clec6a* are significantly upregulated after pulp exposure in *Sh3bp2^+/KI^* mice compared to *Sh3bp2^+/+^* mice; **(B)** No significant differences between *Sh3bp2^+/+^* and *Sh3bp2^+/KI^* mice in *Clec4a2*, *Clec12a*, and *Clec2d* expression. Each dot in graph represents a single biological sample. Data are presented as mean ± SD. Statistical analysis was performed by two-way ANOVA with Sidak test for multiple comparisons. (\**p*<0.05, \*\**p*<0.01, \*\*\*\**p*<0.0001, ns= no significant difference).

### Suppression of *Clec4e* (Mincle) in *Sh3bp2^+/KI^Mincle^KO/KO^* mice

To study the role of Mincle in AP in *Sh3bp2^+/KI^* mice, we crossed *Sh3bp2^+/KI^* mice with *Mincle^KO/KO^* mice to obtain *Sh3bp2^+/+^Mincle^+/+^*, *Sh3bp2^+/KI^Mincle^+/+^*, and *Sh3bp2^+/KI^ Mincle^KO/KO^* mice. The mRNA level of *Clec4e* was mostly abolished in *Sh3bp2^+/KI^ Mincle^KO/KO^* mice (Fig 2A). Although pulp exposure moderately induced Mincle expression in *Sh3bp2^+/KI^Mincle^KO/KO^* mice, the abundance was significantly lower compared to *Sh3bp2^+/KI^Mincle^+/+^* mice (Fig 2A). *Sh3bp2^+/KI^Mincle^KO/KO^* mice showed specific suppression of *Clec4e*, as we observed that changes of *Clec4a2*, *Clec4d*, *Clec6a* mRNA levels before and after pulp exposure were similar to *Sh3bp2^+/KI^Mincle^+/+^* mice (Fig 2B).

**Fig 2:**
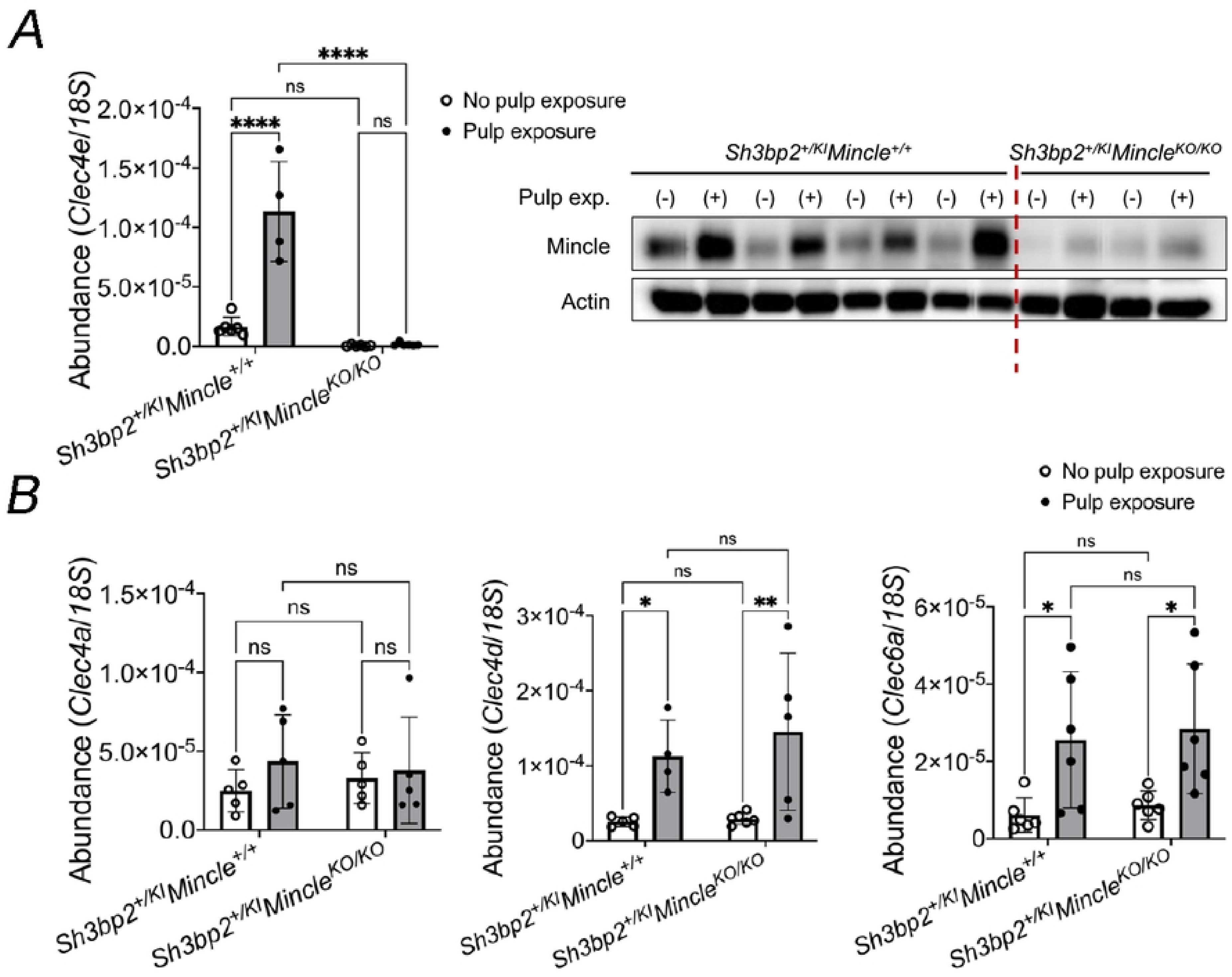
Suppression of Mincle in *Sh3bp2^+/KI^Mincle^KO/KO^* mice. **(A)** *Clec4e* (*Mincle*) mRNA (left panel) and protein (right panel) levels are successfully suppressed in *Sh3bp2^+/KI^Mincle^KO/KO^* mice shown by qPCR and immunoblotting, respectively. Actin serves as sample loading control. Each dot in graphs represents a single biological sample. Statistical analysis was performed by two-way ANOVA with Sidak test for multiple comparisons. (\*\*\*\**p*<0.0001, ns= no significant difference); **(B)** *Clec4a2*, *Clec4d,* and *Clec6a* levels are not altered in *Sh3bp2^+/KI^Mincle^KO/KO^* mice. Each dot in graph represents a single biological sample. Data are presented as mean ± SD. Statistical analysis was performed by two-way ANOVA with Sidak test for multiple comparisons. (\**p*<0.05, \*\**p*<0.01, \*\*\*\**p*<0.0001, ns= no significant difference).

### Comparable mandibular phenotype in Sh3bp2^+/KI^Mincle^+/+^ and Sh3bp2^+/KI^ Mincle^KO/KO^ mice

We monitored body weight changes in *Sh3bp2^+/+^Mincle^+/+^*, *Sh3bp2^+/KI^Mincle^+/+^*, and *Sh3bp2^+/KI^Mincle^KO/KO^* mice days 1-13 after pulp exposure. There were no significant differences between the three groups, suggesting that all mice tolerated the procedures well (Fig 3A). We have previously reported that pulp exposure resulted in jawbone expansion in *Sh3bp2^+/KI^* mice (15). *Sh3bp2^+/KI^Mincle^KO/KO^* mice and *Sh3bp2^+/KI^Mincle^+/+^* mice developed a similar degree of jawbone expansion in contrast to *Sh3bp2^+/+^Mincle^+/+^* mice (Fig 3B). Both *Sh3bp2^+/KI^Mincle^+/+^* and *Sh3bp2^+/KI^Mincle^KO/KO^* mice showed a significantly reduced bone mass after pulp exposure compared to their wild type littermates (Fig 3C).

**Fig 3:**
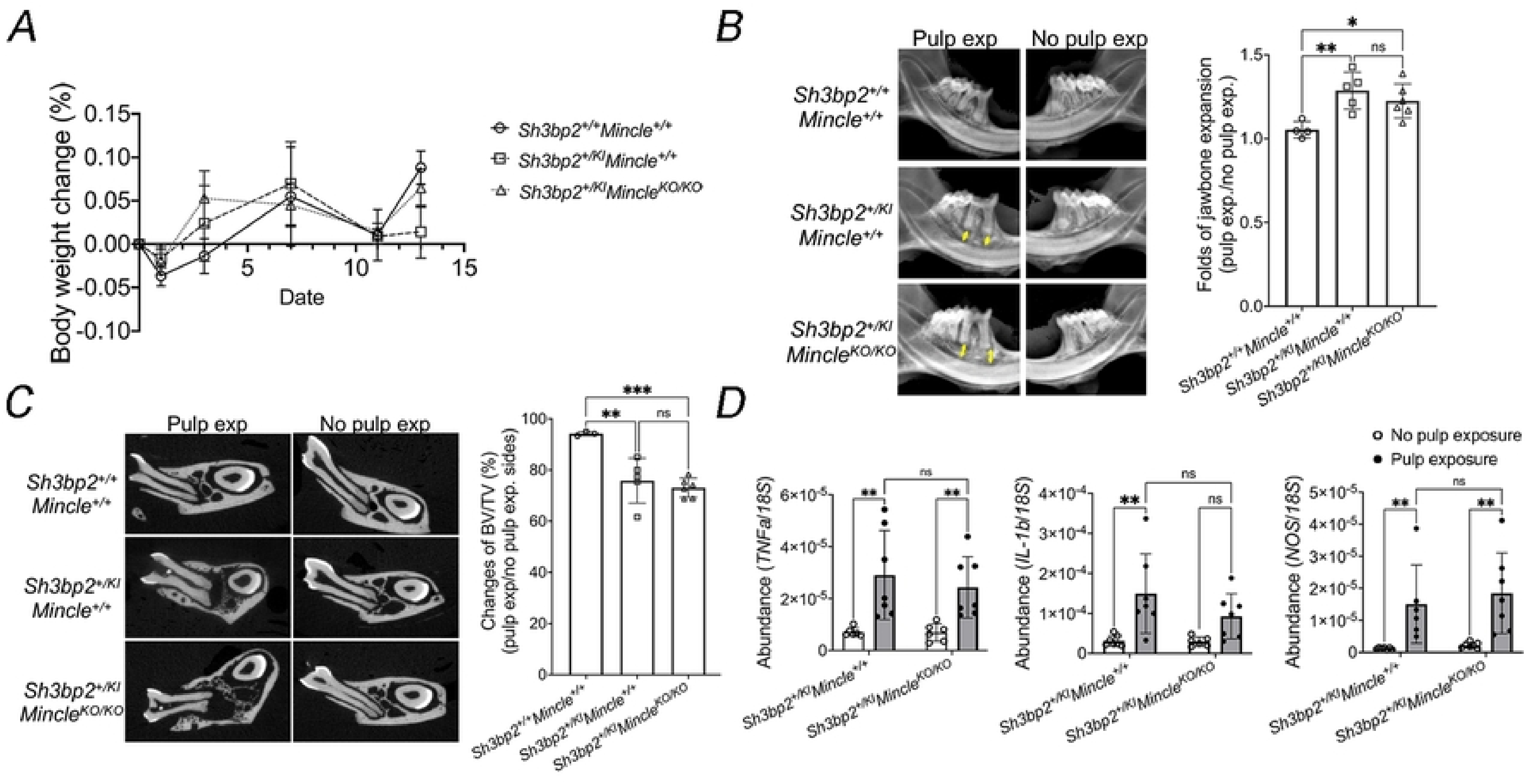
Exacerbated AP in *Sh3bp2^+/KI^Mincle^+/+^* and *Sh3bp2^+/KI^Mincle^KO/KO^* mice. **(A)** Average body weight change ((day_n_–day_n-1_)/day_n-1_x100%) 0-14 days after pulp exposure (n=6-7 per group); **(B**) Representative KUBTEC images showing jawbone expansion (left panel). Quantification is shown on the histogram (right panel). Double headed arrows show increased distance from 1^st^ molar apex to incisors after pulp exposure in *Sh3bp2^+/KI^Mincle^+/+^* and *Sh3bp2^+/KI^Mincle^KO/KO^* mice; **C**) Representative μCT images of mandibles 42 days after pulp exposure (left panel) and percentage changes of BVF (pulp exposure/no pulp exposure mandibles) in *Sh3bp2^+/+^Mincle^+/+^, Sh3bp2^+/KI^Mincle^+/+^,* and *Sh3bp2^+/KI^Mincle^KO/KO^* mice (right panel). Statistical analysis was performed by one-way ANOVA with Tukey’s post-hoc test. (\**p*<0.05, \*\**p*<0.01, \*\*\*\**p*<0.0001, ns= no significant difference). **(D)** Pulp exposure-triggered expression of *TNF-α* (left panel), *IL-1β* (middle panel), and *iNOS* (right panel) in infected mandibles compared to non-infected mandibles in *Sh3bp2^+/KI^Mincle^+/+^* and *Sh3bp2^+/KI^Mincle^KO/KO^* mice by qPCR. Each dot in graph represents a single biological sample. Statistical analyses were performed using two-way ANOVA followed by Sidak correction. (\*\**p*<0.01 indicate statistical significance, ns= no significant difference).

We next examined the induction of inflammatory cytokines by pulp exposure in *Sh3bp2^+/KI^Mincle^+/+^* and *Sh3bp2^+/KI^Mincle^KO/KO^* mice. *IL-1β* and *TNF-α* are the most common inflammatory cytokines detected in periapical lesions (26). Pulp exposure resulted in significantly increased levels of *TNF-α* in *Sh3bp2^+/KI^Mincle^+/+^* and *Sh3bp2^+/KI^Mincle^KO/KO^* mice (Fig 3D). *IL-1β* was significantly increased in *Sh3bp2^+/KI^Mincle^+/+^* mice and showed a tendency of increase in *Sh3bp2^+/KI^Mincle^KO/KO^* mice (Fig 3D). Inducible nitric oxide synthase (*iNOS*) produces high, sustained level of nitric oxide during inflammation (27). Our data show that *iNOS* levels were significantly triggered by pulp exposure in both, *Sh3bp2^+/KI^Mincle^+/+^* and *Sh3bp2^+/KI^Mincle^KO/KO^* mice (Fig 3D). These data demonstrated that suppression of *Clec4e* by a genetic approach does not ameliorate the severity of AP in *Sh3bp2^+/KI^* mice.

### AptMincle partially protects bone loss in AP of *Sh3bp2^+/KI^* mice

We next administrated a highly specific neutralizing DNA aptamer targeting Mincle (3.5mg/kg) to suppress Mincle activation *in vivo*. Affinity and specificity of AptMincle have been validated *in vitro* and an *in vivo* protocol had been established (24). AptMincle was injected into *Sh3bp2^+/KI^* mice starting from the day of pulp exposure, then twice a week for 3 weeks. At the end of 6 weeks, the bone loss specified by the ratio of BVF between pulp exposed and no pulp exposed mandibles, was significantly reduced in *Sh3bp2^+/KI^* mice that received AptMincle injection compared to vehicle treatment (Fig 4). These data demonstrated that short-term suppression of Mincle during the initial stage of inflammation partially rescues the exacerbated AP in *Sh3bp2^+/KI^* mice.

**Fig 4:**
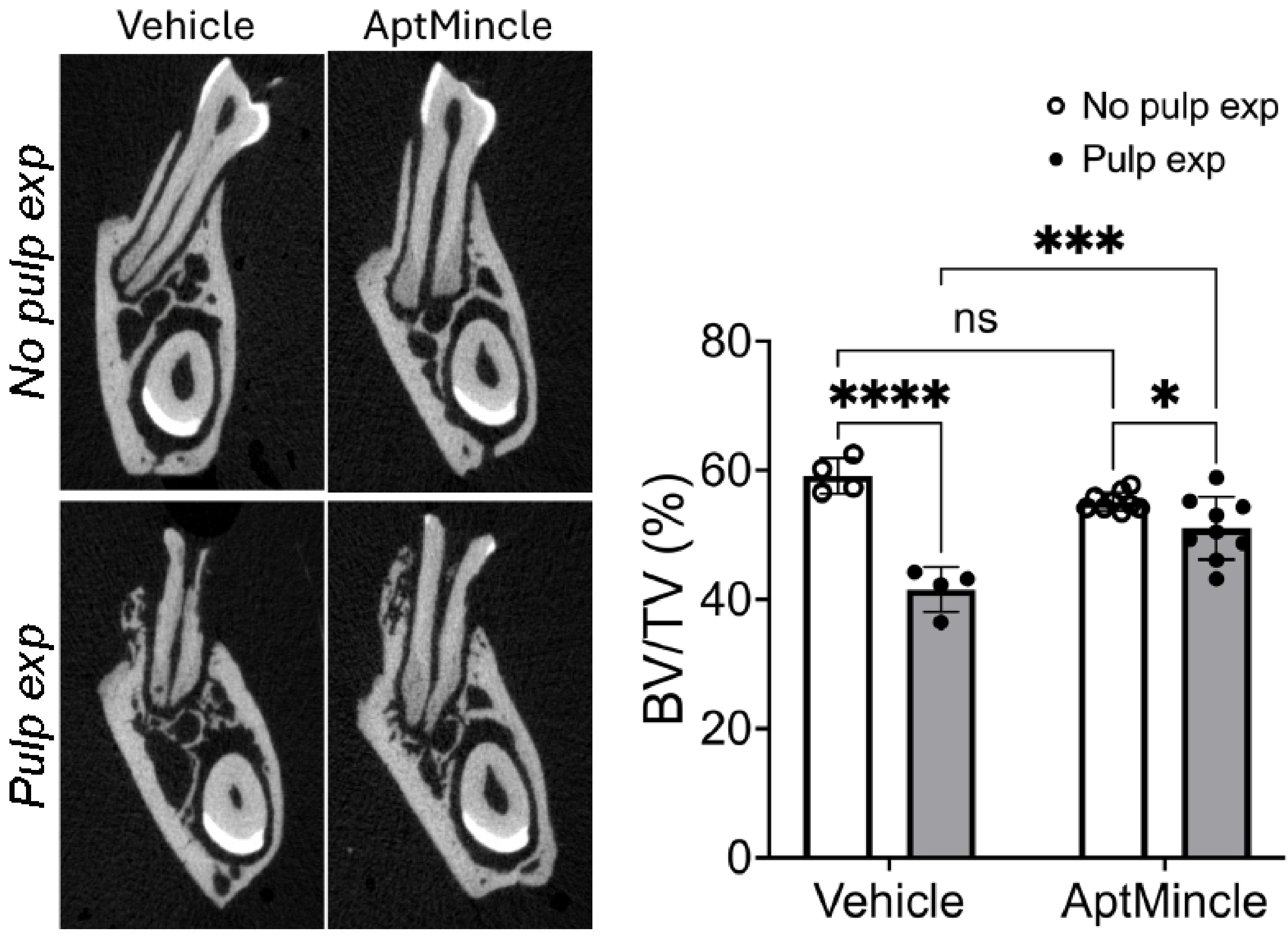
AptMincle ameliorates bone loss in AP in *Sh3bp2^+/KI^* mice. Representative μCT images of pulp exposed and no pulp exposed mandibles from *Sh3bp2^+/KI^* mice treated with vehicle compared to AptMincle. Each dot in graph represents a single biological sample. Data are presented as mean ± S.D. Statistical analyses were performed using two-way ANOVA followed by Sidak correction. (\**p*<0.05, \*\*\**p*<0.0005, \*\*\*\**p*<0.0001 indicate statistical significance, ns= no significant difference).

Taken together, these results confirmed that endodontic infection in *Sh3bp2* mutant mice results in more severe bone destruction compared to wild type mice. Multiple CLRs were triggered by pulp exposure in *Sh3bp2^+/KI^* mice, especially *Clec4e*. Suppression of *Clec4e* by a Mincle KO mouse model is not sufficient to ameliorate bone destruction and block proinflammatory cytokines induced by pulp exposure. Interestingly, the short-term AptMincle injection to reduce Mincle activation successfully provided protective effects for bone loss in *Sh3bp2^+/KI^* mice.

## Discussion

To our knowledge, this is the first study to suggest that hyperactivation of Mincle is at least in part contributing to an increased AP lesion size after pulp exposure in *Sh3bp2^+/KI^* mice. AP is caused by microorganisms and byproducts such as lipopolysaccharide (LPS) following pulp exposure (1). While the oral microflora in humans is more complex than in mice, the immune reaction is generally similar in regards to proinflammatory cytokine expression, bone loss mechanisms, and immune cells involvement in AP (28–30). Oral microbiota are not significantly different between *Sh3bp2^+/+^* and *Sh3bp2^+/KI^* mice (31). Thus, we believe that our results primarily reflect the differential host immune responses between *Sh3bp2^+/+^* and *Sh3bp2^+/KI^* mice.

We suspected that Mincle was highly induced in *Sh3bp2^+/KI^* mice upon PAMPs and DAMPs co-stimulation because: 1) Mincle is mainly expressed in myeloid cells (32) and CBM is primarily driven by hyperactivated myeloid cells (5). 2) Mincle is strongly induced by inflammation, infection, or tissue damage. PAMPs (oral bacteria) and DAMPs during jawbone remodeling are suspected to cause the jaw-specific development of CBM lesions (33). 3) Activated Mincle signals increase inflammatory pathways (NF-κB, MAPK), leading to cytokine production (TNF-α) via FcRγ and Syk kinase. These inflammatory pathways and cytokines are significantly triggered in *Sh3bp2^KI/KI^* mice (12, 14, 34). 4) Mincle is upregulated during early osteoclast differentiation and Mincle activation triggers osteoclastogenesis via ITAM-based calcium signaling pathways (35). Osteoclastogenesis is significantly increased in CBM mice. Our data showed that Mincle was highly induced in *Sh3bp2^+/KI^* mice only after pulp exposure but not in WT littermates, suggesting a functional connection between mutant Sh3bp2 and Mincle proteins. Interestingly, Mincle is induced upon PAMPs-stimulation in *Sh3bp2^+/KI^* but was already high in naïve no pulp exposed *Sh3bp2^KI/KI^* mice (S1 Fig.) suggesting that Mincle may be a determining factor for phenotypic severity in cherubism and can recognize DAMPs as ligands in sterile inflammation in *Sh3bp2^KI/KI^* mice.

Naïve *Sh3bp2^+/KI^* mice are phenotypically closer to *Sh3bp2^+/+^* mice but also developed significantly increased bone loss and jawbone expansion after pulp exposure. In *Sh3bp2^+/KI^Mincle^KO/KO^* mice, Mincle expression was significantly reduced but still developed exacerbated AP with no alteration in expression of *TNFα* and *iNOS* compared to *Sh3bp2^+/KI^Mincle^+/+^* mice. Mincle-deficient mice have been studied previously using different models or stimuli. Mincle-deficient mice exhibited rapid lethality and severe bacteremia upon Group A *Streptococcus* infection, suggesting Mincle provides protective immunity (36). Mincle-deficient mice increase susceptibility to septic injury due to impaired bacterial clearance and neutrophil recruitment (37). Lee et al., concluded that Mincle has a dual role during bacterial infection by activating proinflammatory genes in the promotion of inflammatory responses and is required for subsequent resolution of inflammation in macrophages (38).

To inactivate Mincle at the initial stage of bacterial infection, we adapted an *in vivo* AptMincle protocol that had been used to significantly reduce the severity of ulcerative colitis in mice (24). The rescue effect of colonic inflammation by AptMincle (0.35, 1, or 3.5 mg/kg) was dose-dependent, with 3.5 mg/kg being the most effective. AptMincle injection significantly, although not fully, protected *Sh3bp2^+/KI^* mice from bone loss after pulp exposure (bone loss: 25% vehicle vs. 8% AptMincle). *Sh3bp2^+/KI^ Mincle^KO/KO^* mice failed to rescue the bone phenotype in AP possibly because these mice have impaired clearance of PAMPs/DAMPs, leading to a “frustrated” resolution of inflammation. AptMincle injection rescued the phenotype in *Sh3bp2^+/KI^* mice partly but not fully, possibly due to compensatory mechanisms from other *Clec* genes. Taken together, the continuous inhibition of Mincle in *Sh3bp2^+/KI^Mincle^KO/KO^* mice did not ameliorate bone destruction following endodontic infection. However, inhibition of Mincle activation by injecting *Sh3bp2^+/KI^* mice with AtpMincle at the beginning of the inflammatory stage partially rescued the AP phenotype.

In summary, in this study, we showed that Mincle is hyperactivated in *Sh3bp2^+/KI^* mice after a PAMPs-driven endodontic infection. Our data demonstrated that *Sh3bp2^+/KI^* mice in Mincle-deficient background developed exacerbated AP lesions after pulp exposure, while AptMincle injection provided significant protection from bone loss. Future studies will focus on the crosstalk of *Clec* genes with other PRRs, such as TLRs, and their synergistic or compensatory effects on modulating immune responses during infection and inflammation in CBM. LPS, which is a ligand of TLR2 or TLR4, can activate Mincle (38, 39). Other PRRs, such as NOD1/2 and Mincle have synergistic effects on cytokine production (40). This study also suggests that patients with CBM or other autoimmune diseases may benefit from more frequent dental monitoring and earlier intervention to prevent the progression of endodontic infection, which could exacerbate cherubism lesion development. Regulation of Mincle could be a new concept for ameliorating AP progression in CBM patients.

## Conclusion

Our study showed that Mincle was remarkably induced by endodontic infection in an autoinflammatory disease model for CBM (*Sh3bp2^+/KI^* mice), but not in their WT littermates. Data from our previous and current studies demonstrated that PRRs, including Tlr2/4 and Mincle, are highly induced in the cherubism mouse model, and likely leads to increased infection susceptibility and significantly more alveolar bone destruction in AP. Suppression of Mincle may protect AP progression in this autoinflammatory condition but further research will be needed to better understand Mincle regulation. This study suggests that the patients diagnosed with CBM or other autoinflammatory diseases are likely facing more severe tissue destruction caused by dental infection. Better understanding the mechanisms leading to hyperinflammatory responses is clinically important to improve the quality of dental health for these patients.

## Data Availability Statement

The data that support the findings of this study are available from the corresponding author upon reasonable request.

## Ethics Statement

All animal studies were performed in accordance with animal protocol AP-200505-0424 approved by the Animal Care Committee at the University of Connecticut Health (UConn Health).

## Author Contributions

A.H.: data collection, data analysis and interpretation, performed statistical analyses, revising the paper, final approval to the submitted version. K.W. and E.N.: data collection, data analysis and interpretation, final approval to the submitted version. DF.K. and L.B.: data collection, final approval to the submitted version. Y.U.: study conception and final approval to the submitted version. EJ.R. and A.K.: study conception and design, writing and revising the paper, final approval to the submitted version. IP. C.: study conception and design, data collection, data interpretation, writing and revising the paper, final approval to the submitted version.

## Acknowledgement

The project was supported by InCHIP-SDM, Convergence grants at UConn Health (IP.C.), and R01D025664 from the National Institutes of Health (IP.C.).

